# Comparison of Gene Expression During Early Phases of Limb Regeneration Between Regeneration-permissive Neotenic and Regeneration-deficient Metamorphic Axolotl

**DOI:** 10.1101/693911

**Authors:** Mustafa Sibai, Ebru Altuntaş, Barış Ethem Süzek, Betül Şahin, Cüneyd Parlayan, Gürkan Öztürk, Ahmet Tarık Baykal, Turan Demircan

## Abstract

The axolotl (*Ambystoma Mexicanum*) salamander, an urodele amphibian, has an exceptional regenerative capacity to fully restore an amputated limb throughout the life-long lasting neoteny. By contrast, when axolotls are experimentally induced to metamorphosis, attenuation of the limb’s regenerative competence is noticeable. Here, we sought to discern the proteomic profiles of the early stages of blastema formation of neotenic and metamorphic axolotls after limb amputation by means of LC-MS/MS technology. We quantified a total of 714 proteins having an adjusted p < 0.01 with FC greater or equal to 2 between two conditions. Principal component analysis revealed a conspicuous clustering between neotenic and metamorphic samples at 7 days post-amputation. Different set of proteins was identified as differentially expressed at all of the time points (1, 4, and 7 days post-amputations against day0) for neotenic and metamorphic stages. Although functional enrichment analyses underline the presence of common pathways between regenerative and nonregenerative stages, cell proliferation and its regulation associated pathways, immune system related pathways and muscle tissue and ECM remodeling and degradation pathways were represented at different rate between both stages. To validate the proteomics results and provide evidence for the putative link between immune system activity and regenerative potential, qRT-PCR for selected genes was performed.

## INTRODUCTION

Regeneration, functional restoration of amputated or damaged part of the body, is an important mechanism observed in various cells, tissues and organs of invertebrates and vertebrates at different extent (Brockes et al. 2001; Poss 2010). Regenerative capacity varies greatly in animal kingdom, and within a species, tissues exhibit a diverse range of regeneration ability (Tanaka and Reddien 2011). Although most of the metazoan species somehow experience this process, some invertebrates such as planarians and hydras demonstrate exceptional regenerative skills manifested in the execution of whole body restoration from a small trunk (Holstein et al. 2003; Sánchez Alvarado 2006). On the other hand, for vertebrates, a progressive decline in regenerative capacity with complexity is a commonly accepted phenomena in invertebrates (Gardiner 2005).

Among the tetrapods, urodeles have a remarkable regeneration potential. Axolotl, *Ambystoma Mexicanum*, a member of the urodele group of amphibians, possess spectacular abilities to restore the missing part of the body following an injury. Scar-free wound healing (Denis et al. 2013), entire regeneration of brain (Maden et al. 2013), spinal cord (Mchedlishvili et al. 2007), internal organs (Cano-Martínez et al. 2010) and extremities (Echeverri and Tanaka 2002; Kragl et al. 2009) throughout its life while retaining an astonishingly-low cancer incidence (Ingram 1971; McCusker and Gardiner 2011) are the notable regenerative capability of axolotl. Renewal of complex structures such as entire limb by axolotl, positions it as an attractive model for regeneration research.

Axolotl limb regeneration is accomplished in three main steps: wound healing, neural-epithelial interactions, and reprogramming of positional information (French et al. 1976; Gardiner et al. 1986; Singer 2004). When a limb is amputated, wound healing program is initiated with secretion of pro-regenerative signals that lead to recruitment of the epithelial cells in and around the wound zone to form a permissive wound epithelium (WE) (Gardiner et al. 1986; Endo et al. 2004). Infiltration of immune cells, especially macrophages, is required and indispensable for removal of the pathogens and phagocytosis of dead cell debris from the wound area (Godwin and Brockes 2006; Godwin et al. 2013). Innervation induces dedifferentiation of keratinocytes to gain stem cell like features, a crucial transformation to form apical epithelial cap (AEC) (Satoh et al. 2008). Beneath the WE/AEC, differentiated cells undergo dedifferentiation as a result of nerve dependent gene expression accompanied by secretion of regulatory signals from keratinocytes. Cells reenter the cell cycle and start to accumulate in wound bed. This results in formation of a regenerative specific tissue, blastema, which is a distinctive tissue of epimorphic regeneration that is enriched in population of regeneration-competent progenitor cells (McCusker et al. 2015). While AEC and blastema moves from the proximal towards to distal, cells localized in proximal part start to re-differentiate with growth and patterning signals to form a fully patterned miniature limb that will continue growing (McCusker and Gardiner 2013).

Besides its regenerative capabilities, one of the axolotl’s most interesting features is its life-long neoteny, in other words, retaining the embryonic characters throughout the adulthood since they do not undergo metamorphosis naturally. Metamorphosis defines the biological process by which some animals exhibit a drastic morphological, anatomical and physiological alteration (Brown and Cai 2007) at post embryonic development. For most of the amphibian clade members some time after birth, metamorphosis is observed as a natural mechanism to adapt themselves to terrestrial life conditions. During this process a series of epigenetical changes causes remodeling or disappearing of existing organs and formation of new ones. Although axolotls considered as paedomorphic, metamorphosis can be induced by administration of thyroid hormones (McCusker and Gardiner 2011). For a better understanding of such morphological changes in axolotl, a comprehensive histological atlas of the morphological alterations taking place within tissues and organs throughout metamorphosis has been described (Demircan et al. 2016). In addition to morphological and histological changes, metamorphosis of axolotl brings about a sharp reduction in regeneration rate, and leads to carpal and digit malformations (Monaghan et al. 2014; Demircan et al. 2018). Obtained results from axolotl are well aligned with the previous studies on *Xenopus laevis* which demonstrate dramatic restriction in regenerative capacity with metamorphosis (Mescher and Neff 2006; Godwin and Rosenthal 2014). Furthermore, studies on regeneration-deficient animal models suggest that there is a correlation between cellular changes during ontogenic development and progressive loss of regenerative ability (Seifert et al. 2012; Lee-Liu et al. 2018). Since regenerative capacity differs significantly for developmental stages of amphibians (Mescher and Neff 2006), inducible metamorphosis during adulthood provides a unique experimental advantage to eliminate the impact of age and body size associated changes on regeneration. Therefore, facultative metamorphosis in axolotl offers an ideal system to compare studies between regenerative and nonregenerative forms of Axolotl at the same developmental stage to identify the cellular and molecular mechanism of regeneration process.

Utilization of Axolotl as a fruit model requires generation of reference resources for this animal. Axolotl has a colossal (~32 GB) (Nowoshilow et al. 2018) and complex diploid genome with 14 pairs of chromosome (FANKHAUSER and HUMPHREY 2007). Until very recently (Nowoshilow et al. 2018), obtaining a reference genome for axolotl has been a great challenge due to insufficient read-length, highly repetitive composition of DNA and incompetent methodologies for genome assembly. Recent efforts to elucidate microbial diversity of Axolotl during metamorphosis (Demircan et al. 2018) and limb regeneration (Demircan et al. 2019) has expanded the reference datasets. But still, much of the established knowledge on axolotl limb regeneration has relied on and is mostly still contingent upon, microarray such as (Knapp et al. 2013; Voss et al. 2015) and high-throughput technologies including transcriptomics (e.g.(Bryant et al. 2017; Leigh et al. 2018)) and proteomics (Rao et al. 2009; Demircan et al. 2017). In spite of the valuable insights which RNA based gene expression analysis methods have to offer, an inherent limitation of them lies within their failure to accurately reflect the gene expression activity up to the protein level. Therefore, proteomics holds a great promise to investigate the differential protein expression at different timepoints of regeneration.

Importantly, when referring to axolotl regeneration omics studies, the stage we are referring to is mostly the neotenic stage. To the best of our current knowledge, there have not been omics-based studies aiming at unravelling the limb regenerative cues that distinguish neotenic from metamorphic axolotl limbs. In this study, we, therefore, took the initiative of performing proteomic analysis to identify the proteomic profiles of the regeneration-permissive neotenic axolotl limbs and those of the weakly-regenerative metamorphic axolotl limbs at different timepoints (dpa 0,1,4 and 7) post amputation. 714 differentially expressed proteins were identified by processing of LC-MS/MS data and this protein list is used for further analysis. 82 of the proteins were found as differentially expressed (adjusted p < 0.01, |FC| > 2) in all of the timepoint comparisons (Day1, 4 and 7 against Day0) and a total of 8 commonly-differentially expressed proteins were noticed for both stages, 5 of which had an opposite expression directionality. Gene ontology annotations and KEGG pathway enrichment analyses showed a dramatic decrease in muscle-specific proteins at 7 days post-amputation in both animal stages, however this decline was found sharper in metamorphic samples. Furthermore, at the latter amputation timepoint, neotenic samples appear to have an obstructed activation of the complement and coagulation cascades, unlike metamorphic samples which exhibit a relatively moderate activation of these immunity-related cascades. This perceived correlation between immune system activation, and the competence of regeneration in both animal’s stages was further fostered by qRT-PCR analyses, whereby several proinflammatory cytokines were found significantly up-regulated and down-regulated in metamorphic and neotenic samples, respectively, at 7 days post-amputation compared to 0- and 1-days post-amputation. The obtained results from neotenic and metamorphic axolotl provide novel insights on regulation of limb regeneration and generated proteomics data will contribute to a better understanding of decline in regenerative capacity after metamorphosis.

## RESULTS

In previous studies (French et al. 1976; Monaghan et al. 2014) it was shown that, unlike neotenic axolotls, metamorphic axolotl exhibits delayed limb blastema formation in comparison to neotenic one associated with restricted regeneration capacity. The difference in timing of limb blastema formation among regenerative and non-regenerative stages of axolotls prompted us to study early blastema gene expression profile at the protein level. Proteins at different time points (day0,1,4 and 7) of axolotl limb tissues were identified by using Progenesis QI with a generated protein database for axolotl. In total 714 differentially expressed (fold change greater than or equal to 2.0 between the at least two conditions) proteins were identified with adjusted p value set to p< 0.01 (Figure 1b)

**Figure 1:**
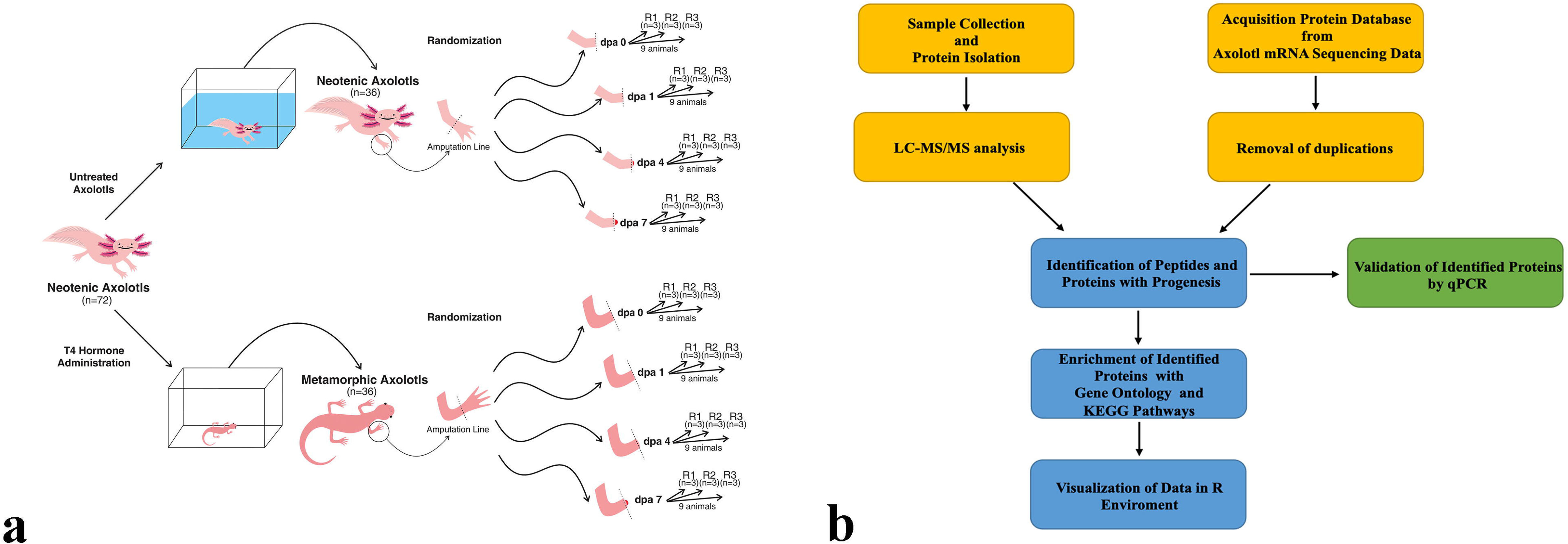
Experimental design and work-flow. a) Generation of the protein datasets in this study is carried out by collecting post amputation limb tissue samples derived from neotenic and metamorphic axolotls. (36 neotenic and 36 metamorphic axolotls limb tissue samples). Samples were obtained from right forelimbs at 0 dpa, 1 dpa, 4 dpa, and 7 dpa; dpa indicates day post amputation. At each experimental time point groups, 9 animals were sub-grouped into 3 biological replicates (R1, R2 and R3) and each replicate was formed by pooling limb tissue samples from 3 animals to minimize variation between animals. b) The workflow of data analysis, the employed tools and changes in the number of processed reads at each step are shown.

### Neotenic and metamorphic samples clustered distantly at later timepoints

Principal Component Analysis (**Figure 2a**) illustrates how metamorphic and neotenic samples cluster together at the early timepoints day0 and day1. On the other hand, a distant clustering between neotenic and metamorphic samples at the later timepoints day4 and day7 was observed, while metamorphic samples cluster together more adjacently compared to how neotenic samples cluster together at both of the later timepoints.

**Figure 2:**
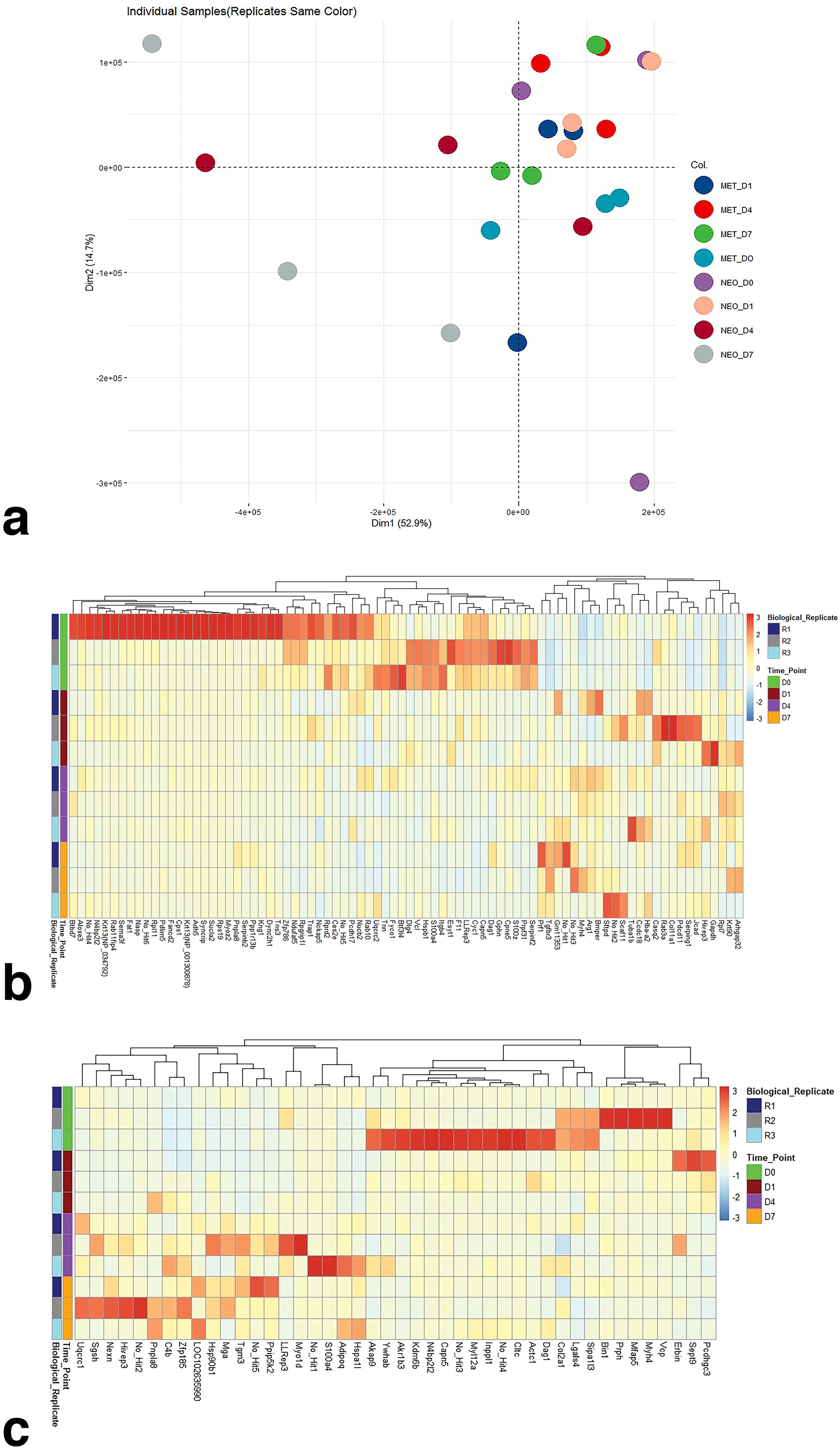
Linear dimensionality reduction and clustering heatmaps of DE proteins. a) Principal Component Analysis (PCA) was performed on DE proteins (adj P <0.01, |FC| > 2) among neotenic and metamorphic samples at 0, 1, 4, and 7 days post-amputation. DE proteins among 0, 1, 4, and 7 days post-amputation in a) neotenic samples) and in b) metamorphic samples were hierarchical clustered using “Complete Linkage” and samples were clustered using “Manhattan” distance. Red asterisks indicate proteins having an opposite expression directionality between neotenic and metamorphic samples at the corresponding amputation timepoint, while green asterisks indicate otherwise.

### Neotenic and metamorphic samples share DE proteins in all timepoints

We performed differential expression analysis for neotenic and metamorphic samples to identify the proteins that had altered expression levels in response to limb amputation. The number of proteins which were found differentially expressed (adjusted p < 0.01, |FC| > 2) in all of the timepoint comparisons day1/day0, day4/day0, and day7/day0 were 82 DE proteins for neotenic samples (**Figure 2b**) and 44 DE proteins for metamorphic samples (**Figure 2c**). A total of 8 proteins were detected in both neotenic and metamorphic DE protein lists, out of which 5 proteins included calcium-independent phospholipase A2-gamma (*Pnpla8*), metastasin (*S100a4*), 40S ribosomal protein S2 (*LLRep3*), myosin heavy chain-4 (*Myh4*) and human immunodeficiency virus type I enhancer-binding protein 3 (*Hivep3*) had an opposite differential expression directionality while the other 3 proteins; NEDD4 binding protein 2-like 2 (*N4bp2l2*), Calpein 5 (*Capn5*), and Dystroglycan precursor (*Dag1*) had a similar DE directionality.

### Various GO terms were enriched by DE proteins in neotenic and metamorphic samples at day7/day0

To understand the early time of blastema formation in neotenic and metamorphic axolotls we focused on DE proteins of 7 dpa samples. GO enrichment analysis was separately carried out on the lists of up-regulated and down-regulated proteins of neotenic and metamorphic samples at day7/day0.

A total of 25 biological processes (BP), 27 cellular components (CC), and 15 molecular functions (MF) were enriched by the 68 up-regulated proteins (**Supplementary Table 1**) in neotenic samples at day7/day0, while the other 136 down-regulated proteins enriched a total of 38 BPs, 68 CCs, and 21 MFs (**Supplementary Table 2**). The 152 up-regulated proteins in metamorphic samples at day7/day0 enriched a total of 53 BPs, 44 CCs, and 28 MFs (**Supplementary Table 3**), whereas the other 144 down-regulated proteins enriched a total of 76 BPs, 78 CCs, and 26 MFs (**Supplementary Table 4**).

Processes such as myosin filament assembly and organization, skeletal muscle thin filament assembly, skeletal myofibril assembly, sarcomere and intermediate filament cytoskeleton organization, cardiac muscle fiber development and tissue morphogenesis, and collagen metabolic process were the top 10 BPs enriched by the up-regulated proteins in neotenic samples at day7/day0 (**Figure 3a**), while negative regulation of hemostasis, actin filament organization, exocytosis and its regulation, protein activation cascade, response to metal ion, fibrinolysis, chondrocyte development, regulation of heterotypic cell-cell adhesion, and negative regulation of coagulation were the top 10 BPs enriched by the up-regulated proteins in metamorphic samples at day7/day0 (**Figure 3c**). Those up-regulated proteins in neotenic samples at day7/day0 also enriched cellular components, such as A band, intermediate filament and cytoskeleton, sarcomere, myosin 2 complex, contractile fiber, myofibril, M band, and actin cytoskeleton (**Figure 3a**). On the other hand, the up-regulated proteins in metamorphic samples enriched similar and other cellular components, such as collagen-containing extracellular matrix, actin cytoskeleton, myelin sheath, contractile fiber part, basement membrane, myofibril, contractile fiber, intermediate filament, and cell cortex (**Figure 3c**). Molecular functions such as structural constituent of cytoskeleton, muscle alpha-actinin binding, structural constituent of muscle, actinin and ankyrin binding, and endopeptidase inhibitor activity were among the top 10 MFs enriched by the top up-regulated proteins in neotenic samples at day7/day0 (**Figure 3a**), however extracellular matrix structural constituent, coenzyme binding, carbohydrate binding, phospholipid binding, calcium transporting ATPase activity, glycosaminoglycan binding, and coupled, ATPase activity were among the top 10 MFs enriched by the up-regulated proteins in metamorphic samples at day7/day0 (**Figure 3c**). Other molecular functions such as actin and actin filament binding, as well as calcium-dependent phospholipid binding were among the top 10 MFs and commonly enriched by the up-regulated proteins in neotenic and metamorphic samples at day7/day0 (**Figure 3a,c**).

**Figure 3:**
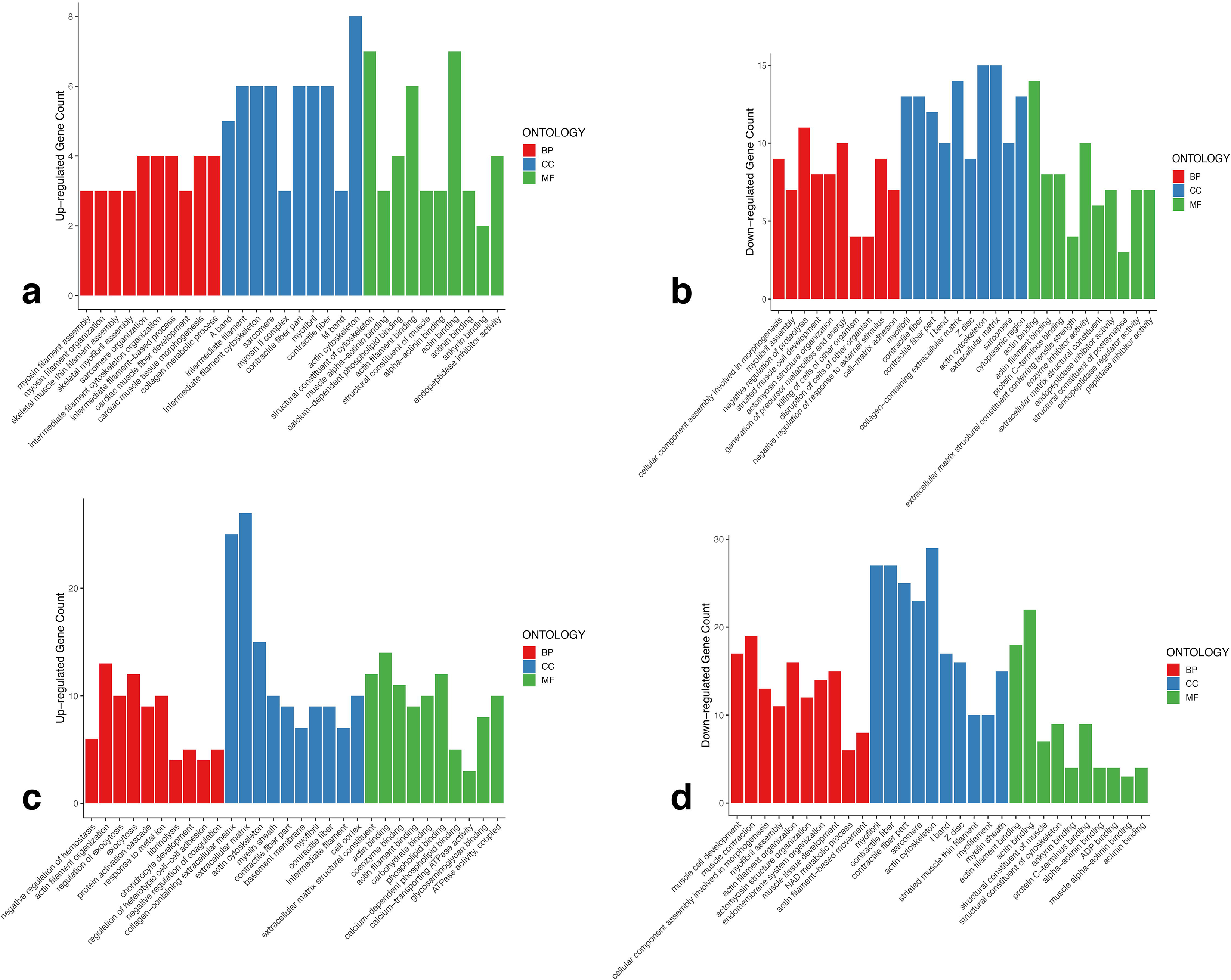
Gene ontology enrichment by DE proteins in each animal stage at 7 days compared to 0 days post-amputation. The top 10 enriched terms (adj P < 0.05) of each GO category (BP, CC, MF) by their corresponding DE protein (adj P < 0.01, |FC| > 2) count were plotted using barplots. a,b) GO terms enriched by up-regulated proteins and down-regulated proteins, respectively, in neotenic samples. c,d) GO terms enriched by up-regulated proteins and down-regulated proteins, respectively, in metamorphic samples.

Among the top 10 BPs enriched by the down-regulated proteins in neotenic samples at day7/day0 were negative regulation of proteolysis, striated muscle cell development, generation of precursor metabolites and energy, killing and disruption of cells of other organism, negative regulation of response to external stimulus, and cell-matrix adhesion (**Figure 3b**). On the other hand, other BPs such as muscle cell development and contraction, actin filament organization, endomembrane system organization, muscle tissue development, NAD metabolic process, and actin filament-based movement were among the top 10 BPs enriched by the down-regulated proteins in metamorphic samples at day7/day0 (**Figure 3d**). BPs such as cellular component assembly involved in morphogenesis, myofibril assembly, and actomyosin structure organization, as well as CCs such as myofibril, contractile fiber, sarcomere, actin cytoskeleton, I band, and Z disc were among the top 10 BPs and top 10 CCs and commonly enriched by the down-regulated proteins in neotenic and metamorphic samples at day7/day0 (**Figure 3b,d**). Collagen-containing extracellular matrix and cytoplasmic region were among the top 10 CCs enriched by the down-regulated proteins in neotenic samples at day7/day0 (**Figure 3b**), while striated muscle thin filament, myofilament, and myelin sheath were among the top 10 CCs enriched by the down-regulated proteins in metamorphic samples at day7/day0 (**Figure 3d**). The three molecular functions actin binding, acting filament binding, and protein C-termins binding were among the top 10 MFs and commonly enriched by the down-regulated proteins in neotenic and metamorphic samples at day7/day0 (**Figure 3b,d**). Other top MFs enriched by the down-regulated proteins in neotenic samples at day7/day0 were extracellular matrix structural constituent conferring tensile strength, enzyme inhibitor activity, endopeptidase inhibitor and regulator activity, peptidase inhibitor activity, and structural constituent of postsynapse (**Figure 3b**), whereas structural constituent of muscle and cytoskeleton, ankyrin binding, alpha-actinin binding, ADP binding, and actinin binding were among the top 10 MFs enriched by the down-regulated proteins in metamorphic samples at day7/day0 (**Figure 3d**).

### Various KEGG terms were enriched by DE proteins in neotenic and metamorphic samples at Day7/Day0

KEGG enrichment analysis was carried out on the lists of DE proteins (up-regulated and down-regulated ones combined) of neotenic and metamorphic samples at Day7/Day0. A total of 4 KEGG pathways were enriched by the DE proteins in neotenic samples at day7/day0; estrogen signaling pathway, dilated cardiomyopathy, hypertrophic cardiomyopathy, and complement and coagulation cascades (**Figure 4a**). Those four pathways were also found among the 23 pathways enriched by the DE proteins in metamorphic samples at day7/day0 (**Figure 4b**). The other enriched pathways including the carbon metabolism, pyruvate metabolism, ECM-receptor, focal adhesion, cardiac muscle contraction are listed in **Figure 4b**. Thereafter, the coagulation and complement cascades were further investigated due to their relevance to immunity processes which may be importantly unique in identifying the distinctive regenerative patterns in neotenic and metamorphic samples.

**Figure 4:**
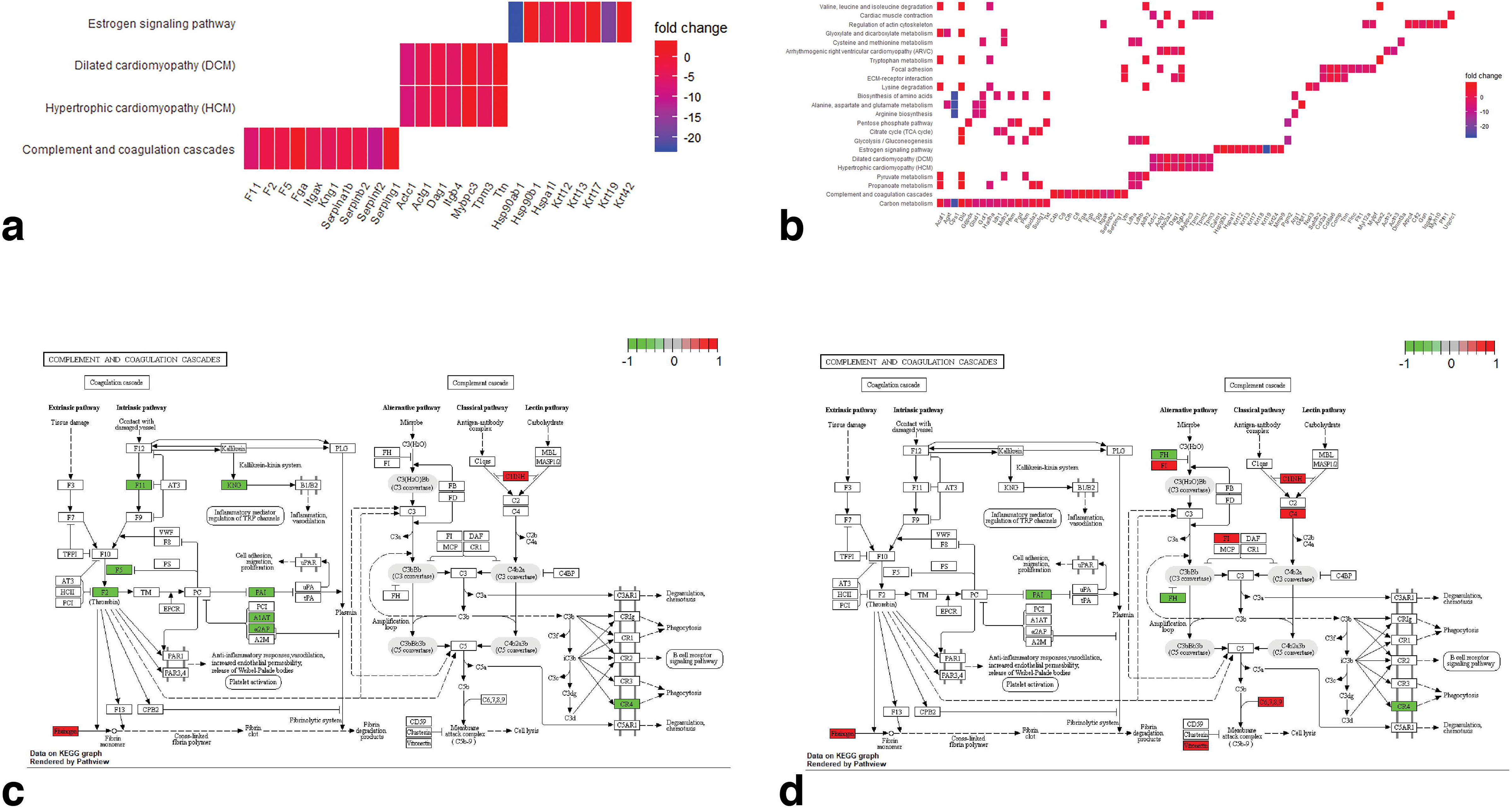
KEGG pathway enrichment by DE proteins in each animal stage at 7 days compared to 0 days post-amputation. All KEGG enriched pathways (adj P < 0.05) by DE proteins (adj P <0.01, |FC| > 2) in a) neotenic and b) metamorphic samples were visualized using heatplots. The DE proteins enriched in complement and coagulation cascades in c) neotenic and c) metamorphic samples were rendered and visualized as part of the whole two cascades on a KEGG graph. Considering all DE proteins have a |fold change| > 2, the colored bar of the KEGG graphs is a qualitative representation of the fold change whereby a scale of 1 and above (red) refers to a status of up-regulation and a scale of −1 and below refers to a status of downregulation.

The coagulation cascade proteins included coagulation factor XI (F11), coagulation factor V (F5), prothrombin (F2), kininogen (KNG), plasminogen activator inhibitor (PAI), α-1-antityrpsin (A1AT), and α-2-antiplasmin (α2AP) were found down-regulated in neotenic samples at day7/day0 (**Figure 4c**), however only PAI protein in this cascade in metamorphic samples was found down-regulated at day7/day0 (**Figure 4d**). Fibrinogen was found up-regulated in the coagulation cascade in both neotenic and metamorphic samples at day7/day0 (**Figure 4c,d**). As for the complement cascade, an up-regulated plasma protease C1 inhibitor (C1INH) and a down-regulated CR4 were found in both neotenic and metamorphic samples at Day7/Day0 (**Figure 4c,d**). Furthermore, among the complement cascade components, FI, C4, C6, C7, C8, C9, vitronectin proteins were up-regulated and FH protein was down-regulated in metamorphic samples at day7/day0 (**Figure 4d**). In order to have a deeper understanding of the differential protein expression of the complement and coagulation cascades in neotenic and metamorphic samples at Day7/Day0, each set of up-regulated and down-regulated proteins were separately tested for KEGG enrichment in neotenic and metamorphic samples, respectively.

Out of the total 5 KEGG pathways enriched by the down-regulated proteins in neotenic samples at day7/day0, hypertrophic cardiomyopathy and dilated cardiomyopathy were also found enriched by the down-regulated proteins in metamorphic samples at day7/day0 (**Supplementary Tables 5 and 6**). However, importantly, the complement and coagulation cascades were found enriched by the up-regulated and down-regulated proteins at day7/day0 in metamorphic samples and neotenic samples, respectively (**Supplementary Tables 5 and 7**).

### Neotenic and metamorphic samples shared DE proteins at day7/day0, some of which have opposite expression directionality

The 204 DE proteins in neotenic samples at day7/day0 were merged with the 296 DE proteins in metamorphic samples at day7/day0. A total of 130 genes were found common between the two sample types at this timepoint comparison. In order to test whether those common genes all have the same, different, or a mix of protein expression directionality, a spearman correlation test was conducted on their fold changes. Spearman’s coefficient was reported as 0.39, indicating a slightly-low but positive correlation. Out of those 130 proteins, 47 had an opposite direction of protein expression (**Figure 5a**).

**Figure 5:**
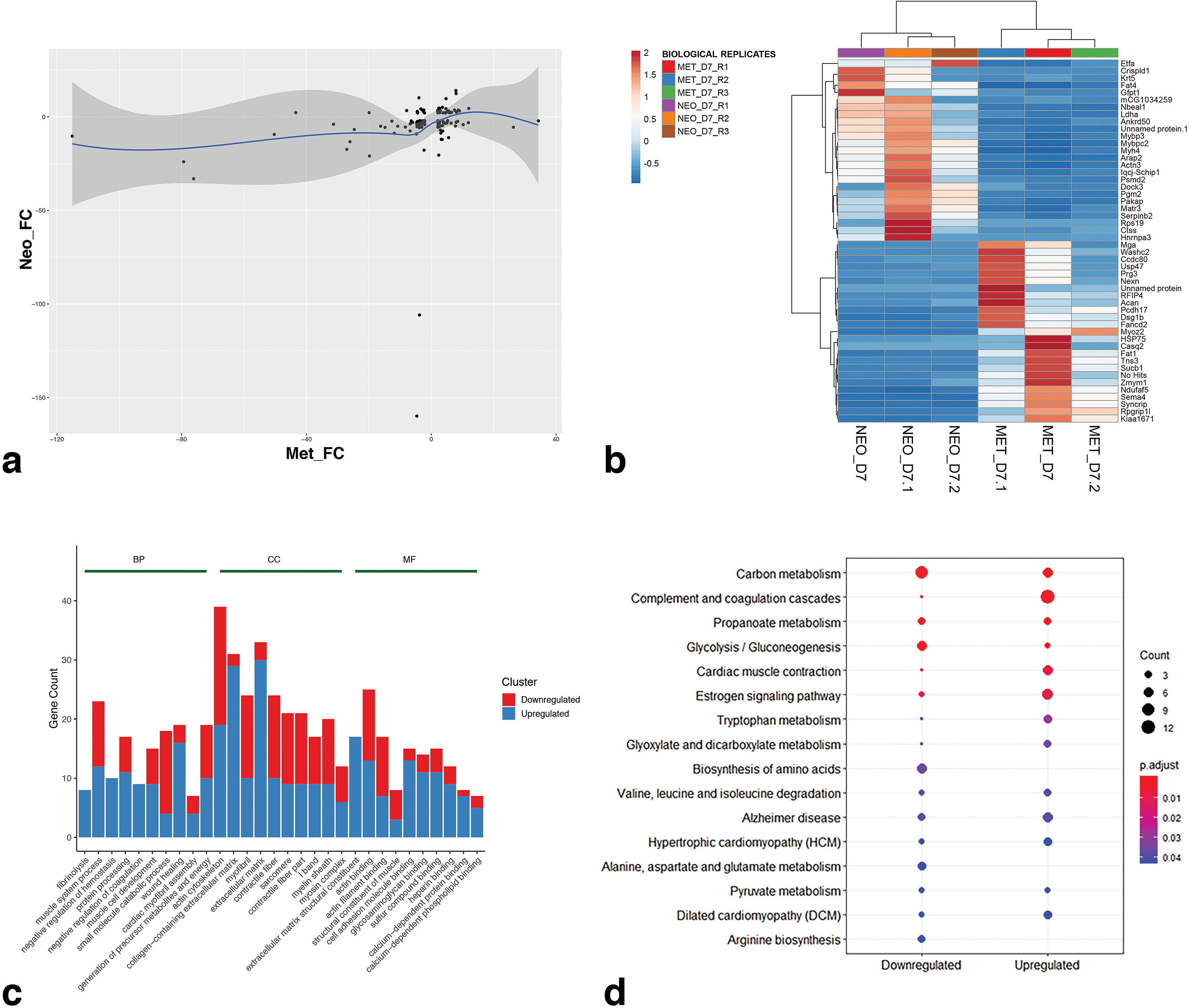
Correlation testing, clustering, and GO and KEGG enrichment. a) Scatter plot showing the fold changes of DE proteins at day7 compared to day0 post-amputation shared in metamorphic samples (x-axis) and neotenic samples (y-axis). b) Correlation distance and Average linkage-based clustering heatmap was used to visualize the top 50 DE (25 up-regulated, 25 down-regulated) proteins (adj P < 0.01, |FC| > 2) at day 7 post-amputation in metamorphic samples compared to neotenic samples. c) Barplots were used to visualize the top 10 enriched terms (adj P < 0.05) of each GO category (BP, CC, MF) by their corresponding DE protein (adj P < 0.01, |FC| > 2) count in metamorphic samples compared to neotenic samples at day 7 post-amputation. d) All KEGG enriched pathways (adj P < 0.05) by DE proteins (adj P <0.01, |FC| > 2) in metamorphic samples compared to neotenic samples at day 7 post-amputation were visualized as clusters of up-regulated and down-regulated proteins using dotplot.

**Figure 6:**
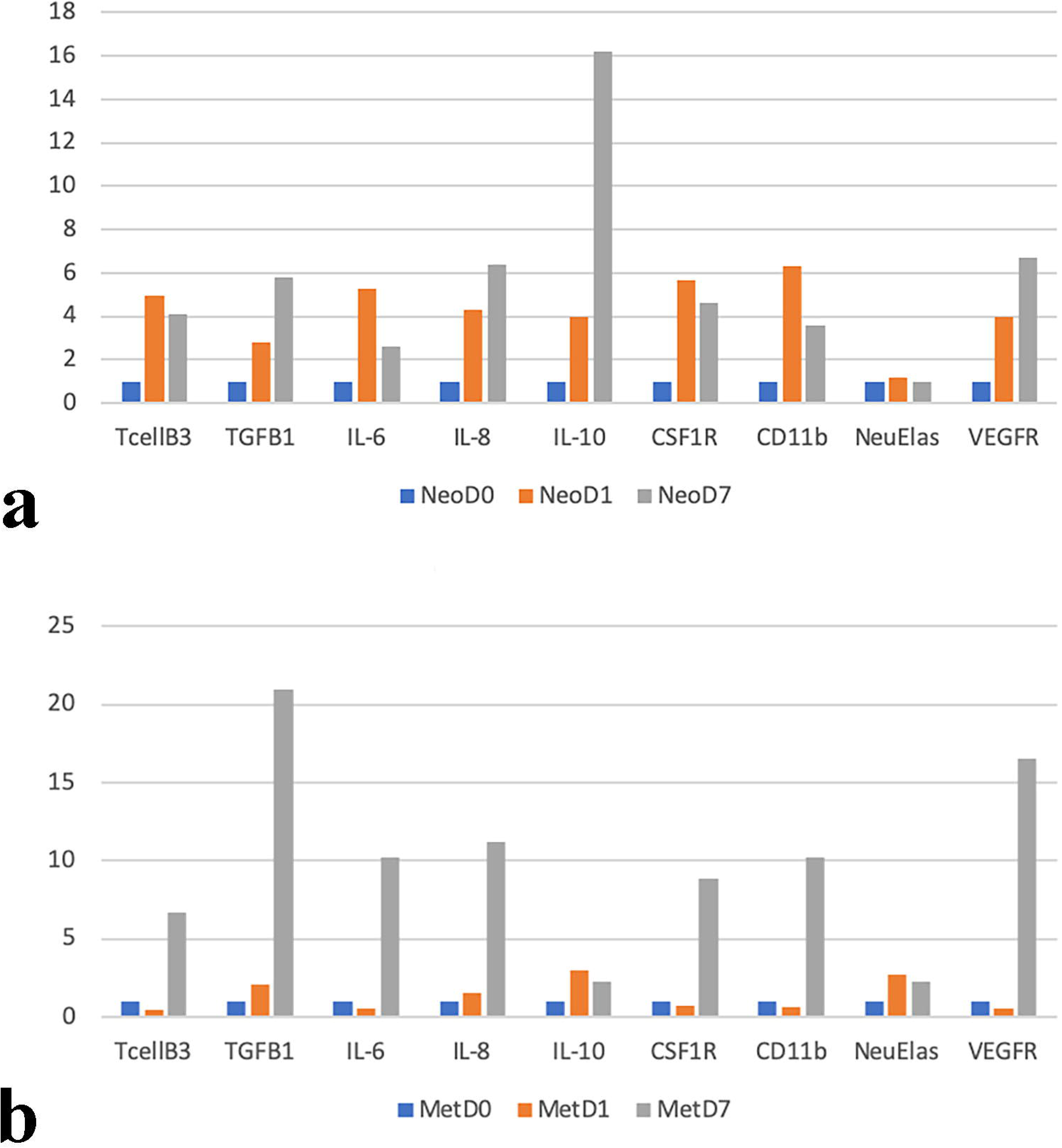
qRT-PCR of selected immune-modulating genes for a) neotenic and b) metamorphic samples at 1 and 7 dpa compared to day0. TGFB1: transforming growth factor beta 1, IL-6: Interleukin 6, IL-8: Interleukin 8, IL-10: Interleukin 10, CSF1R: colony stimulating factor 1 receptor, CD11b: cluster of differentiation molecule 11B, NeuElas:Neutrophil elastase, VEGFR: Vascular Endothelial Growth Factor Receptor

### Clustering and Enrichment of the DE proteins in metamorphic day7 samples versus neotenic day7 samples

A total of 259 proteins were differentially expressed at 7 dpa in metamorphic samples compared to neotenic samples, out of which 163 were up-regulated (FC > 2) and 96 were down-regulated (FC > 2). The 25 most up-regulated proteins such as aggrecan (*Acan*), tensin-3(*Tns*), T-cell-mediated immunity related Semaphorin 4A (*Sema4*), and the 25 most-downregulated proteins included myosin heavy chain 4 (*Myh4*), endosome-to-plasma membrane trafficking related ankyrin repeat domain-containing protein 50 (*Ankrd50*), dedicator of cytokineses 3 (Dock3) associated with axonal outgrowth in metamorphic Day7 samples to neotenic Day7 samples were represented on a heatmap (**Figure 5b**).

In order to complement the information gathered from the resultant processes and pathways which were observed in neotenic and metamorphic samples at day7/day0, GO and KEGG enrichment analyses were also carried out on the 259 DE proteins in metamorphic samples with respect to neotenic samples at day7 timepoint. Those DE proteins enriched a total of 139 BPs, 92 CCs, 60 MFs, and 16 KEGG pathways (**Supplementary Tables 8 and 9**).

Among the top 10 BPs enriched by the DE proteins in metamorphic day7/neotenic day7 samples (**Figure 5c**), fibrinolysis, negative regulation of hemostasis, and negative regulation of coagulation were enriched completely by up-regulated proteins. Protein processing, muscle cell development, and wound healing were enriched by mostly up-regulated proteins. Muscle system process, cardiac myofibril assembly, and generation of precursor metabolites and energy were enriched by an almost equal amount of up-regulated and down-regulated proteins. Small molecule catabolic process was enriched by mostly down-regulated proteins. As for the top 10 CCs (**Figure 5c**), actin cytoskeleton, I band, and myosin complex were enriched by an almost equal amount of up-regulated and down-regulated proteins. collagen-containing extracellular matrix and extracellular matrix were enriched by mostly up-regulated proteins. Myofibril, contractile fiber, sarcomere, and myelin sheath were enriched by mostly down-regulated proteins. Regarding the top 10 MFs (**Figure 5c**), extracellular matrix structural constituent was enriched completely by up-regulated proteins. Actin binding was enriched by an almost equal amount of up-regulated and down-regulated proteins. Structural constituent of muscle and actin filament binding were enriched by mostly down-regulated proteins. Cell adhesion molecule binding, glycosaminoglycan binding, sulfur compound binding, heparin binding, calcium-dependent protein binding, and calcium-dependent phospholipid binding were enriched by mostly up-regulated proteins.

The four KEGG pathways estrogen signaling, complement and coagulation cascades, HCM, and DCM, which were previously found enriched in metamorphic and neotenic samples at day7/day0 timepoint (**Figure 4a,b**), were all enriched mostly by up-regulated proteins in metamorphic samples with respect to neotenic samples at day7 timepoint (**Figure 5d**). Other pathways which were mostly enriched by up-regulated proteins were cardiac muscle contraction, tryptophan metabolism, and glyoxylate and dicarboxylate metabolism. Carbon metabolism and glycolysis/gluconeogenesis were mostly enriched by down-regulated proteins. Biosynthesis of amino acids, arginine biosynthesis, and alanine, aspartate and glutamate metabolism were completely enriched by down-regulated proteins. Pathways which were enriched by an equal or almost equal proportions of up-regulated and down-regulated proteins were propanoate metabolism, pyruvate metabolism, and valine, leucine and isoleucine degradation. Notably, all of the latter 11 pathways (**Figure 5d**) were also enriched by the DE proteins in metamorphic samples at day7/day0 (**Figure 4b**). However, Alzheimer disease was the single pathway enriched in metamorphic samples with respect to neotenic samples at 7 dpa (**Figure 5d**) that was not observed in any of the enrichment results in neotenic and metamorphic samples of day7/day0 timepoints. The complement and coagulation cascades were also further investigated. The coagulation cascade proteins F11, F2, A1AT, α2AP, A2M, and fibrinogen were found up-regulated at 7 dpa in metamorphic samples compared to neotenic samples (**Supplementary Figure XY**). The complement cascades, on the other hand, was enriched by the down-regulated protein FH and the up-regulated proteins C1INH, C5, C6, C7, C8, C9, and vitronectin at 7 dpa in metamorphic samples compared to neotenic ones.

### qRT-PCR Analysis of the Expression Profile of Immunity Markers at Early Stages of Blastema Formation in Neotenic and Metamorphic Axolotls

We performed qRT-PCR in neotenic and metamorphic samples at 0,1, and 7 dpa to understand the immune responses that occur as a result of amputation and during blastema formation (**Figure ??**). The expression levels of T cell receptor (TcellB3) and colony stimulating factor 1 receptor (CSF1R) in neotenic samples were upregulated at day 1 compared to day 0 and then they showed a downtrend at day 7 with respect to day 1. Metamorphic samples didn’t exhibit significant changes in terms of expression level of TcellB3 and CSF1R at day 0 and day 1, however they were dramatically upregulated at day 7. To gain insight about pro-inflammatory responses we examined the expression level of Interleukin-6 (IL-6) and Interleukin-8 (IL-8). In neotenic samples, IL-6 presented an upregulation at day 1 compared to day 0 and a downregulation at day 7 compared to day 1. IL-8 had similar expression profile with IL-6 at day 1 in neotenic samples; however, unlike IL-6, continued upregulation of IL-8 was observed at day 7. At day 1, metamorphic samples exhibited downregulation of IL-6 and upregulation of IL-8 compared to day 0. In addition, IL-6 and IL-8 expression levels were found dramatically upregulated at day 7 in metamorphic samples. Interleukin-10 (IL-10), an anti-inflammatory cytokine, showed a remarkable alteration in neotenic samples. It presented an upregulation in day 1 samples compared to day 0 samples. The main finding of IL-10 is that its expression level is strikingly increased at day 7 with respect to day 0 and day 1. Metamorphic samples, on the other hand, had a negative correlation with neotenic samples in terms of IL-10 expression level. Even though IL-10 had an uptrend at day 1, its expression level decreased at day 7. We also analyzed the expression profiles of transforming growth factor beta 1 (TGFB1) and vascular endothelial growth factor receptor (VEGFR) in neotenic and metamorphic samples. TGFB1 and VEGFR had similar expression profiles in neotenic samples, in which they were upregulated at 1 and 7 dpa compared to 0 dpa. When examining the expression levels of TGFB1 and VEGFR in metamorphic samples, although they were similarly upregulated at day 7 with respect to day 1 and day 0, TGFB1 was upregulated and VEGFR was downregulated at day 1 compared to day 0.

## DISCUSSION

While animals such as zebrafish, lizards, and urodeles are able to regenerate some of their extremities even after reaching adulthood, frogs and mammals lose this ability prior to metamorphosis (Seifert et al. 2012) or during post embryonic development. Although neotenic axolotls have an exceptional regenerative capacity, previous studies (Demircan et al. 2018, 2019) have shown defective and infidel limb regeneration of axolotl after metamorphosis. Since findings of published studies underlined an apparent delay in phases of regeneration for metamorphic animals, in this study we aimed to reveal the gene expression pattern at early phases of regeneration (day0,1,4 and 7) for both neotenic and metamorphic axolotls at the protein level.

Principle component analysis revealed a clear clustering between metamorphic and neotenic samples at Day 7 post amputation, suggesting overall biological alterations taking place distinctly between the animal stages at the limb regenerates at this timepoint. Although one can expect that Day 0, Day1, and Day4 post amputation samples should also cluster distantly, this was not observed in our PCA which is likely due the fact that the identified proteins are mostly common between these samples with a similar expression level.

Testing for the differential expression of proteins at all four timepoints at once between neotenic and metamorphic samples revealed that each of the three proteins which had similar DE directionality (Dag1, Capn5, and N4bp2l2) contribute to the regenerative program in both animal stages alike. Dystrophin associated glycoproteins (Dag1), a cytoskeleton-linked extracellular matrix receptor, plays a role in negatively regulating cell migration (Ibraghimov-Beskrovnaya et al. 1992; Brennan et al. 2004), and since it is down-regulated at 1,4, and 7 dpa in both animal stages, this allows cell migration events to take place via the inhibition of cell-cell adhesion. Calcium-dependent cysteine protease 5 (Capn5), the vertebrate homolog of *Caenorhabditis elegans* TRA3 which is known for mediating necrotic pathways in neurons (Barnes and Hodgkin 1996), was down-regulated at 1,4, and 7 dpa in both animal stages, which is known that blockage of necrotic event in neurons is required for formation of AEC and a successful limb regeneration (Salley and Tassava 1981). NEDD4-binding protein 2-like 2 (N4bp2l2), also known as PFAAP5, was also found as down-regulated at 1,4, and 7 dpa in both animal stages. PFAAP5 interacts with immature neutrophil elastase (ELA2) and growth factor independent-1 (Gfi1) (Salipante et al. 2009). It has been postulated that PFAAP5 plays an essential role in mediating interactions between ELA2 and Gfi1 to contribute to neutrophil differentiation or neutropenia (Salipante et al. 2009). In a previous study it has been demonstrated that ELA2 levels increase gradually at wound healing stage and returning to low levels at later stages, around 10 days post amputation (Godwin et al. 2013) which is crucial for successful regeneration. Down regulation of PFAAP5 may allow maturation of ELA2 which would be followed by secretion of mature neutrophil elastase to the wound area.

On the other hand, five proteins (Hivep3, RPS2, s100a4, Pnpla8, Myh4) which are differentially expressed at all time points within neotenic and metamorphic samples exhibit an opposite or fluctuated gene expression profile. Human immunodeficiency virus type 1 enhancer binding protein 3 (Hivep3) can induce the expression of T-cells’ Interleukin 2 (IL2) (Oukka et al. 2004). IL2 promotes T regulatory (Treg) cells’ development that act as immunosuppressive cells, suppressing effector T cells’ proliferation (Bettelli et al. 2006). Tregs are particularly important for timely wound-reepithelization (Nosbaum et al. 2016). The up-regulation of Hivep3 at 1 dpa in neotenic samples, may promote wound re-epithelialization through IL2-mediated activation of Tregs. Metamorphic samples, however, up-regulate hivep3 at around 7dpa, indicating a possible delayed activation of wound healing processes. It has been previously hypothesized that successful regeneration may require weak adaptive immunity (Mescher and Neff 2005). Interleukin 4 (IL-4) is important for B and T cells activation (Vallé et al. 1989; Silva-Filho et al. 2014). The LLRep3, also known as 40S ribosomal protein s2 (RPS2), can be induced by the expression of IL4 (Kafasla et al. 2014), and since LLRep3 is down-regulated at 1,4,and 7dpa in neotenic samples, this may indicate an under-expression of IL4, and thus, weak activation of adaptive immunity from 1 to 7 dpa. On the contrary, metamorphic samples up-regulate LLRep3 at 4 dpa, implying an over-expression of IL4 at this timepoint, suggesting a time at which an activation of adaptive immunity may be hindering or delaying the formation of blastema cells. At 1,4, and 7 dpa, S100 calcium binding protein a4 (s100a4) protein was down-regulated for neotenic samples. In metamorphic samples for the same protein, down-regulation at day1 and 7 is accompanied by slight up-regulation at 4 dpa. This protein has an important role in cancer metastasis and is overexpressed in a variety of tumors (Orre et al. 2013). This observation is in-line with the notion of low-cancer incidence feature of axolotls. Although cancer metastasis is known to activate epithelial-to-mesenchymal transition (EMT) program which promotes cell migration, axolotls may still activate this program while maintaining normal cell growth and proliferation. Pnpla8, calcium-independent phospholipase A2-gamma takes role in membrane remodeling and lipid mediator signaling which can be considered as a part of cellular lipid metabolism (Yoda et al. 2010). Pnpla8 acts on membrane phospholipids and it releases free arachidonic acid (AA) which is catalyzed further by cyclooxygenase (COX) isoenzymes to produce active prostaglandins and in return an inflammation response is generated. Neotenic samples down-regulate pnpla8 at 1, 4, and 7 dpa, which may imply an incremental inhibition of strong inflammation response possibly important for the activation of dedifferentiation, cell migration and proliferation processes. Metamorphic samples, however, strongly up-regulate pnpla8 at all time points, which may cause a significant inflammation response and consequently, low rates of cell migration and proliferation may result in delaying blastema formation. Myosin heavy chain-4 (Myh4) was found down-regulated in metamorphic samples at 1,4, and 7 dpa, that might be indication of early and progressive muscle degradation during metamorphic axolotl limb regeneration. Surprisingly though, our results confirm that neotenic samples up-regulate myh4 at those time points which may be perceived as re-organization of muscles instead of rapid degradation.

In the light of these results we decided to focus on 7 dpa, the time at which the major differences during blastema formation between neotenic and metamorphic stages start to occur.

Several GO terms enriched by DE proteins in both neotenic and metamorphic samples at day7/day0, and even when relatively comparing metamorphic to neotenic samples at 7dpa, were part of the coagulation and complement cascades, such as fibrinolysis, negative regulation of coagulation, negative regulation of hemostasis, and plasminogen activity. We, therefore, wanted to take a deeper look at the key proteins and how they are involved in those two cascades by using the kegg pathway viewer, as they may form pivotal roles in the distinctive regenerative features between the two animal stages. The complement and coagulation cascades appear to be obstructed in neotenic samples at day7/day0. Several proteins are down-regulated downstream of the coagulation intrinsic pathway which can lead to anti-coagulation. While an up-regulated fibrinogen is observed (normally required for the formation of fibrin clots), the down-regulation of PAI protein allows the activation of uPA and tPA complex to initiate the production of plasmin, and its direct inhibition is blocked by the down-regulation of α2AP proteins. The plasmin anti-degradation mechanism is further actuated by the down-regulation of F2 (thrombin), rendering a weak conversion of fibrinogen to fibrin monomers, an inactivation of F13 whose activity is required for the conversion of fibrin monomers to cross-lined polymers, as well as the inactivation of CPB2 that is now incapable of blocking the plasmin. Consequently, the fibrinolytic system is activated and fibrin degradation takes place. Concomitantly, the complement system is probably barely activated with the production of plasmin, by which C5 may get activated and membrane attack complex can be generated leading to cell lysis. The cross-talk between fibrinolytic system and the complement activation may suggest why neotenic axolotls are regeneration-permissive in which fibrin clots may act as obstacles towards the formation of new cells at regenerating tip of the limb, and the need for other cells to be lysed at the same time. On the other hand, metamorphic samples at day7/day0 have a much less activated fibrinolytic system, and a relatively moderate expression of plasmin. Any activation of the latter would probably still be able to activate C5, and with C6, C7, C8, and C9 being up-regulated, a definite formation of membrane attack complex may occur, leading to cell lysis. It may be conceivable, therefore, that at 7 dpa metamorphic axolotls start to exhibit regeneration-deficient symptoms at the molecular and cellular level due to the incomplete activation of the fibrinolytic system and the relatively-strong activation of the complement system which may be a response by which cells may sense a cease in proliferation because of the presence of fibrin clots, hence they tend to undergo more lysis. This notion can be further asserted when comparing metamorphic samples to neotenic samples at Day7 timepoint. Indeed, the relative up-regulation of F2, α2AP, and A2M inhibit plasmin from fully activating the fibrinolytic degradation system, and the relative up-regulation of C5 (naturally and by scarce plasmin) along with C6,7,8,9 leads to further cell lysis through the formation of membrane attack complexes.

In neotenic day7/day 0 samples, the vast majority of the top 10 biological processes along with other significant BPs enriched by up-regulated proteins center around muscle tissues. Notably, though, many of them were enriched by very few proteins (each BP having a max of 4 or 5 proteins), most of which were found in almost every process. Myosin filament assembly, myosin filament organization, skeletal muscle thin filament assembly, skeletal myofibril assembly, cardiac muscle fiber development, and muscle fiber development were enriched by the same three proteins (Ttn, Mybpc1, Mybpc3). Another two top processes: intermediate filament cytoskeleton organization and intermediate filament-based process were both enriched by the same set of proteins (Krt17, Nefm, Ina, Vim). In a previous study (Rao et al. 2009), it was demonstrated that upregulation of some proteins associated with muscle contraction may be correlated with mitosis on the basis of chromosome condensation and segregation. Other muscle-related processes were enriched by the same former three proteins along with a couple more per process (Krt8, Tgfbr3, Chga, Fat4, Myh1). On the other hand, only three muscle-related processes were enriched by down-regulated proteins in neotenic day7/day0 samples. Myofibril assembly, striated muscle cell development, and actomyosin structure organization share seven down-regulated proteins (Neb, Actn2, Myom1, Actg1, Actc1, Krt19, Casq2). The latter two processes had an extra Pdlim5 and Serpinf2 down-regulated proteins, respectively. Despite having a lower number of muscle-related BPs enriched by down-regulated proteins than those enriched by up-regulated proteins, those latter 3 processes were among the top 10 BPs and they shared a higher number of proteins. **Therefore, at day7 timepoint, there appears to be a somewhat moderate muscle activity through up-regulation of some proteins and a tendency for an initiation of further muscle degradation through a highly-connected set of down-regulated muscle-related proteins.** Unlike neotenic samples, metamorphic samples at day7/day0 have much less muscle-related biological processes enriched by up-regulated proteins compared to those enriched by down-regulated proteins. Actin filament organization (Hsp90b1, Cap1, Pfn1, Capg, Cit, Fer, Arpc4, Mical3, Cfl2, Pls3, Mybpc1, Fat1, Trf), muscle cell migration (Fga, Adipoq, Vtn, Iqgap1, Anxa1), regulation of smooth muscle cell migration (Fga, Adipoq, Vtn, Iqgap1) and regulation of actin polymerization or depolymerization (Pfn1, Capg, Cit, Fer, Arpc4, Cfl2) were the only four muscle-related BPs, the first of which being amongst the top 10, and were all enriched by up-regulated proteins. Notably, the proteins of the third and fourth processes are also among those in the second and first processes, respectively. On the other hand, a plethora of muscle-related BPs were enriched in metamorphic day7/day0 samples by down-regulated proteins, constituting the vast majority of the top 10 BPs, with the top six of muscle processes sharing a set of five proteins (Mybpc3, Actc1, Mybpc2, Myom1, Actn2), and at least thirty other proteins distributed among those processes. Metamorphic samples at 7dpa hence clearly undergo a dramatic decrease in muscle tissues as demonstrated by the high number of down-regulated proteins in muscle tissues along with the number of processes they enrich. KEGG enrichment analysis provided another layer of evidence to the behavior of muscle-related genes in neotenic and metamorphic samples at day7/day0. Indeed, both of dilated cardiomyopathy (DCM) and hypertrophic cardiomyopathy (HCM) processes were enriched by the same set of 7 proteins in neotenic samples, two of which are up-regulated (Ttn, Mybpc3) and the other five are down-regulated (Dag1, Actg1, Tpm3, Itgb4, Actc1). Those two processes were also enriched in metamorphic samples by the same set of 9 proteins, two of which are up-regulated (Atp2a2, Itgb4) and the other 7 are down-regulated (Actg1, Dag1, Tpm1, Tpm2, Mybpc3, Tpm3, Actc1). This is indicative of the observation that both neotenic and metamorphic axolotls undergo muscle tissue histolysis over the course of limb regeneration. Notably, Dag1, Actg1, Tpm3, Actc1, Itgb4, and Mybpc3 are common between neotenic and metamorphic day7/day0 samples, and the latter two proteins have an opposite expression direction. This observation, where some muscle-specific genes are oppositely differentially expressed, may further assert the notion of different time-shifting in muscle tissue depletion between neotenic and metamorphic axolotls. Based on the current literature, axolotls initiate muscle-tissue degradation at around 1-2 dpa (Dwaraka et al. 2018) and an incremental increase in muscle-tissue histolysis gets accelerated at around 7 dpa until an absolute reduction of muscle tissue is observed at around 10 dpa (Voss et al. 2015). Our data validated the literature for metamorphic axolotls, as they exhibit a sharp decrease of muscle tissues at 7 dpa compared to neotenic ones. Another interesting observation is an overall higher number of up-regulated and down-regulated muscle-related proteins in metamorphic samples than in neotenic ones at day7/day0. In fact, a recent study (Dwaraka et al. 2018) on comparative transcriptomics of limb regeneration among two paedomorphic Ambystoma species (A.Mexicanum and A. Andersoni) and one metamorphic Ambystoma (A.macalatum) detected a steeper down-regulation of muscle-specific genes at 24 hour post amputation (hpa) in metamorphic species compared to the paedomorphic (neotenic) ones. The authors argue that dramatic tissue remodeling in Ambysotma is a characteristic of its ancestral developmental mode, that is metamorphosis, during which the alteration of muscle structure and composition may take place at early stages just before animals transition from the short-lived aquatic larval habitats into terrestrial locomotion. Although the authors derive their hypothesis from their observation of having sharper levels of down-regulated proteins, which we do not quite see in such patterned way in our data, probably due to amputation timepoint differences, we could still relate to it from our observation of having an overall higher number of up-regulated and down-regulated proteins in metamorphic samples compared to neotenic ones.

When comparing metamorphic to neotenic samples at 7 dpa, Keratin 5 (Krt5) was found relatively up-regulated in neotenic samples. Up-regulation of Krt5 was shown to be an important regulator in regenerating blastema (Moriyasu et al. 2012), while its down-regulation is observed in differentiating limbs. Expression levels of aggrecan (Acan) is also increased during cell differentiation (Takagi et al. 2008), which was found down-regulated in neotenic at 7 dpa compared to metamorphic samples. At 7 dpa, neotenic samples are still undergoing cellular dedifferentiation and regeneration and they are yet to transition to re-differentiation which takes place at later timepoints. The relative up-regulation of Krt5 and the down-regulation of Acan in neotenic relative to metamorphic samples also may explain why metamorphic axolotls exhibit a delay in blastema formation. Another protein that is deemed important for cellular differentiation, particularly epidermal keratinocytes’ differentiation, is plasminogen activator inhibitor 2 (Serpinb2) (Jang et al. 2010). Serpinb2 could be down-regulated over the phase of dedifferentiation until the re-differentiation phase commences. Although it seems that neotenic samples over-express serpinb2 relative to metamorphic samples at 7dpa, this protein is actually down-regulated in both animal stages at day7/day0. Notably though, it is much less down-regulated in neotenic than in metamorphic samples which fits with the notion of delayed and obstructed limb regeneration in metamorphic axolotl.

FAT atypical cadherin 1 (Fat1) is relatively up-regulated in metamorphic compared to neotenic samples at 7 dpa. As an upstream protein, Fat 1 takes role in assembly and activation of Hippo pathway (Martin et al. 2018), a key kinase tumor suppressor cascade that regulates cell growth, proliferation, differentiation and organ size (Yu and Guan 2013). As another layer of supporting data, Fat 1 protein in neotenic samples is down-regulated (FC of d7/d0 is ~ −3.58) in regeneration time course, whereas a significant up-regulation of Fat 1 in metamorphic samples (FC of d7/d0 is ~ 2.16) was observed. This may explain the difference in cell division rate of neotenic and metamorphic axolotl limb regeneration shown in a previous published study (Monaghan et al. 2014). On the other hand, FAT atypical cadherin 4 (Fat4), a tumor suppressor gene acting on the Wnt/β-catenin signalling (Cai et al. 2015), is found relatively up-regulated at day7 in neotenic samples compared to metamorphic ones. Another essential function of Fat4 is maintenance of planar cell polarity (PCP) (Zhang et al. 2016), which blastema cells require for exhibiting their positional information. In a recent study (Albors et al. 2015) it has been demonstrated that re-expression of PCP genes is crucial for axolotl spinal cord regeneration by orienting mitotic spindles along to anterior-posterior axis. Fat4 might act on PCP pathway to provide apical-basal and anterior-posterior positioning of blastema cells for neotenic axolotl to support successful regeneration.

Compared to metamorphic samples at 7 dpa, neotenic samples relatively up-regulate cathepsin s (Ctss), a lysosomal enzyme that cleaves some extracellular matrix proteins (Fonović and Turk 2014), they down-regulate coiled-coil domain-containing protein 80 (Ccdc80) and desmoglein 1 beta precursor (Dsg1b), both of which promote cell adhesion (Pulkkinen et al. 2003; Pei et al. 2018), and they also down-regulated Tensin3 (Tns3), which inhibits cell migration and matrix invasion (Martuszewska et al. 2009). Furthermore, down-regulated Ctss and up-regulated Ccdc80, Tns3, and Dsg1b were observed in metamorphic samples at 7dpa/0dpa, and all of which were conversely regulated at 7dpa/0dpa in neotenic samples, further asserting the idea of deferral of EMT activation and blastema formation in metamorphic axolotls. Cleavage of ECM components and loss of cell-cell adhesion, is vital for EMT activation, cell migration, and therefore essential for successful regeneration. Based on our results we can speculate that ECM remodeling is achieved fruitfully in neotenic axolotl rather than the metamorphic group.

Metamorphic axolotls seem to undergo a boost in the activation of adaptive immune system along with heightened inflammatory activities. They up-regulate Semaphorin 4 (Sema4) compared to neotenic samples at 7dpa. Sema4 is an important protein in immune cell trafficking, B cell development, T cell maturation and cytokine production (Xue et al. 2016; Alto and Terman 2017). Overall, Sema4 have a positive role on immune cell migration. Moreover, Proteoglycan 3 (Prg3) is up-regulated in metamorphic samples compared to neotenic samples at 7dpa. It is also up-regulated in metamorphic and down-regulated in neotenic samples at 7dpa/0dpa. Over-expression of Prg3 can induce the release of interleukin 8 (IL-8), a pro-inflammatory chemokine that triggers chemotaxis of neutrophils (Plager et al. 1999), thereby causing immune cell migration towards to the site of regeneration. In a nutshell, metamorphic axolotls seem to have a more active immune system with stronger inflammation response which may obstruct regeneration, as well as hinderance of cell migration.

The striking distinction in immunity profile between neotenic and metamorphic samples gave rise to thought of how the immune system can have a role on successful and deficient regeneration. To find a clue about this argument, by performing qPCR, we analyzed expression profile of some pro and anti-inflammatory cytokines and cell receptors whose roles are well-known in immunity. The qPCR results provide another layer of evidence as to how the immune system may dramatically affect the regenerative potential in both animal stages. The T-cell receptor (TcellB3), Interleukin 6 (IL-6), Interleukin 8 (IL-8), Colony stimulating factor 1 receptor (CSF1R), Cd11b, and vascular endothelial growth factor receptor (VEGFR) serve as reliable readout markers for immune system activity and are known for their role in maturation of immune cells and modulating the activation of the immune system with pro-inflammatory actions. Those immunity markers are upregulated at day1 compared to day0 which aligns well with wound healing phase of neotenic axolotl regeneration. Expression level of these genes either down-regulated or moderately up-regulated at day7 compared to day1 and this considerable modulation of immune profile may facilitate the blastema formation. For metamorphic axolotls, on the other hand, down-regulation or moderately up-regulation of expression level is observed at day1 compared to day0 which is a sign of limited activation of immune system at wound healing phase of regeneration in metamorphic animals. Significant up-regulation of these markers at day7 compared to day0 and day1 indicates a clear shift in immune response timing which might interfere with blastema formation due to excessive pro inflammatory signaling between immune system elements.

Interleukin 10 (IL-10), an anti-inflammatory cytokine, is up-regulated at day1 in both animal stages compared to day0 which might be important to balance the pro-inflammatory signals. IL-10 level is detected as considerably up-regulated at day7 in neotenic axolotls compared to day1, a notable evidence of negative regulation of immune response, whereas, remarkably, decline of IL-10 level at day7 in comparison to day1 implies persistency of immune system activity at day7 in metamorphic samples. Transforming growth factor beta 1 (TGFB1) is commonly considered as an anti-inflammatory cytokine, however recent studies suggest that it may have pro-inflammatory activity under different circumstances (Han et al. 2012). Our qPCR results display up-regulation of TGFB1 at day1 and day7 compared to day0 in both neotenic and metamorphic axolotls. Taken together, the observed profile of expressions of pro- and anti-inflammatory cytokines and markers to immune cell maturation signify a negative correlation between the persistent immune system activity and regeneration capacity as observed in metamorphic axolotls, while in neotenic animals mild and early time activity of the immune system appears to be aiding towards the regenerative program.

## CONCLUSION

Due to the anatomical similarity between the limbs of axolotls and those of humans, understanding the differences in regenerative potential between neotenic and metamorphic axolotls would be conducive to regenerative biology and medicine. Presented data here is the first comparison of *Ambystoma Mexicanum* limb tissues for regeneration-permissive and regeneration-deficient stages at proteome level. Our analyses provide new clues to clarify the biological processes which facilitate or hinder axolotl limb regeneration capacity before and after metamorphosis. Notably, for many of the identified proteins longitudinal gene expression trend was overlapping during neotenic and metamorphic axolotl limb regeneration. This may emphasize the fact that regeneration gene expression program is partly preserved after the metamorphosis. Yet for many other proteins, an opposite direction in expression level is detected between the regeneration time points of both stages. ECM degradation, muscle re-organization and degradation, cell division, differentiation and migration pathways are regulated differentially during axolotl limb regeneration at neotenic and metamorphic stages. In addition to these specified pathways, strikingly, expression level of immune system associated genes differs greatly for neotenic and metamorphic animals throughout the regeneration period. Most probably mild immune response and temporary notion of immune system activity play a central role to facilitate regeneration as well as being active at mainly early stages of this process. To test this hypothesis, gain and loss of function studies should be performed to explore the precise roles of immune system in limb regeneration of axolotl prior to and post metamorphosis.

## MATERIALS AND METHODS

### Experimental Design, Thyroxine-Induced Metamorphosis and Sample Collection

Axolotls were obtained from the Ambystoma Genetic Stock Center and grown and bred at the Istanbul Medipol University Medical Research Center. Animals were housed in %40 Holfreter’s solution and fed once a day using a staple food (JBL Novo LotlM, Neuhofen, Germany). Neotenic and metamorphic axolotls were maintained as one animal per aquarium on a 12:12 light-dark cycle at a constant temperature (18-20 °C). The local ethics committee of Istanbul Medipol University approved all animal protocols and experimental procedures described in this article with authorization number 38828770-E15936.

The experimental design followed in this study is shown in Fig.1a. A total of 72 adult wild-type axolotls (12-15 cm in length, 1 year old) were randomly chosen among the siblings. Out of 72 axolotls, 36 were kept in neoteny and the other 36 were induced to metamorphosis using L-thyroxine (Sigma-Aldrich, St Louis, MO, USA, Cat. No. T2376) as described in (Demircan et al. 2018). Briefly, T4 solution was prepared by dissolving L-thyroxine in Holtfreter’s solution equivalent to 50 nM final concentration. The rearing solution of axolotls was replaced with freshly prepared T4 containing solution every third day. Animals were treated with T4 solution for 6 weeks, after which weight loss, as well as the disappearance of the fin and gills take place. Before the amputation and sample collection steps, metamorphic axolotls were allowed to adapt to terrestrial life conditions for a month without hormone treatment.

The 36 neotenic animals were randomly categorized into four groups representing timepoint amputations (0 dpa, 1 dpa, 4 dpa, and 7 dpa). In each group, 9 animals were sub-grouped into three biological replicates (R1, R2, and R3) to obtain repeatable accuracy. The same grouping was carried out for metamorphic axolotls. All animals were anesthetized prior to amputations using 0.1% Ethyl 3-aminobenzoate methanesulfonate (MS-222, Sigma-Aldrich, St Louis, MO, USA). Amputations were performed on the right forelimb of each animal at mid-zeugopod level. Samples were collected according to the defined timepoint amputations; 0 dpa, 1 dpa, 4 dpa and 7 dpa. To minimize the variation between animals, tissue samples of the three animals making a biological replicate were pooled together. Sample collection at 0 dpa and 1 dpa was performed by cutting approximately 1-mm tissue around the amputation site. Sample collection at dpa4 and dpa7 was carried out by cutting the newly formed blastema and at 0.5 mm posterior tissue from the amputation site. All tissue samples were cryopreserved in liquid nitrogen after the collection and stored at −80°C until qRT-PCR and proteomic analyses.

### Protein extraction and sample preparation

Sample preparation for LC-MS/MS is based on previously published protocols (Gurel et al. 2018). UPX protein extraction buffer (Expedeon) was used for protein extraction as per the manufacturer’s instructions. Using a mini disposable micropestle, the samples were homogenized mechanically. 200 ul UPX buffer was added to each sample and the samples were sonicated in 0.5 second bursts at %50 power for 1 minute using a vial tweeter (Hielscher UP200St) and placed in a 100°C water bath for 5 minutes. After homogenization, the insoluble fractions were removed by centrifugation at 14.000 rpm for 10 minutes. Tryptic digest was obtained by performing filter aided sample preparation (FASP) method (Wiśniewski et al. 2009). Briefly, 50 ug of protein lysate was incubated with dithiothreitol (DTT) and iodoacetamide (IAA) respectively for reduction and alkylation steps. Protein concentration was determined by Bradford Protein Assay prior to trypsinization step. Trypsin (Promega) was then added at 1:50 (w / w) and digestion was carried out for 18 h. Peptide concentration was measured by Quantitative Fluorometric Peptide Assay (Pierce) prior to LC-MS/MS analysis.

### Label-free quantitative nano-LC-MS/MS proteomics analysis

LC-MS/MS-based differential protein expression analysis was performed as in our previous studies (Demircan et al. 2017). In brief, 200 ng tryptic peptide mixture was analyzed by nano-LC-MS/MS system (Acquity UPLC M-Class and SYNAPT G2-si HDMS; Waters. Milford, MA, USA). Before the analyses, columns were equilibrated with 97% mobile phase A (0.1% Formic Acid (FA) in LC-MS-grade water (Merck)) and the column temperature was set to 45 °C. The mass spectrometer was calibrated with a MS/MS spectrum of [Glu1]-Fibrinopeptide B human (Glu-Fib) solution (100 fmol/uL) delivered through the reference sprayer of the NanoLockSpray source.

The peptide samples were separated from the trap column (Symmetry C18 5μm,180μm i.d. × 20 mm) onto the analytic column (CSH C18, 1.7 μm, 75 μm i.d. × 250 mm) by a linear 2-h gradient (4% to 40% Acetonitrile 0.1% (v/v) FA, 0,300 ul/min flow rate). 100 fmol/ul Glu-q-fibrinopeptide-B was used as lock mass reference at 0,500 ul/min flow rate with 60 s intervals. Positive ionization mode was used at 50-2000 m/z in the full scan mode.

Acquisition parameters are used from previously published method (Demircan et al. 2017). Data independent acquisition mode (DIA) was applied to the MS and MS/MS scans using 10 V (low collision energy) and 30 V (high collision energy), respectively. Cycle time was 1.4. Also, the ions were separated according to their drift-time by ion mobility separation (IMS). A wave velocity ramp from 1000 m/s to 550 m/s was applied over the full IMS cycle. Release time for mobilty trapping was set as 500 μs and trap height of 15 V with mobility extract height of 0 V. IMS delay was set to 1000 μs after trap release, for the mobility separation. All the ions within 50-2000 m/z range were fragmented without any precursor ion preselection in resolution mode.

### LC-MS/MS data processing

Progenesis QI for proteomics (v.4.0, Waters) was used for the quantitative analysis of peptide features and protein identification. Analysis of the data included retention time alignment to a reference sample, normalization considering all proteins and peptide analysis. Processing parameters were 150 counts for the low energy threshold and 30 counts for the elevated energy threshold. The principle of the search algorithm has been described in detail previously (Distler et al. 2014). All mass data acquired were imported to Progenesis QI and data analysis was performed using the following parameters: minimum number of fragmented ion matches per peptide = 3, minimum number of fragment ion matches per protein = 7, minimum number of unique peptides per protein = 2, maximum number of one missed cleavage for tryptic digestion, fixed modification = carbamidomethyl C, variable modifications = oxidation M and deamidation N and Q, false discovery rate (FDR) ≤ 1%. Only features comprising charges of 2+ and 3+ were selected. Since there is no Axolotl protein database, previously assembled Axolotl mRNA sequencing data was used to generate a protein database. The sample sets were normalized, based on the total ion intensity. Expressional changes and *p* values were calculated with the statistical package included in Progenesis QI for proteomics and protein normalization was performed according to the relative quantitation using non-conflicting peptides. The resulting data set was filtered by ANOVA p-value ≤ 0.01 and only proteins with a differential expression level between the two conditions greater than or equal to 2.0-fold change were considered.

### Quantitative Reverse Transcription PCR (qRT-PCR)

Total RNA isolation from frozen tissues that were collected at 0, 1, 4, and 7 dpa was performed using TRIzol Reagent (Invitrogen™, Cat. No. 15596018) according to the manufacturer’s protocol. Complementary DNA (cDNA) was synthesized from total RNA of all different time points with M-MuLV Reverse Transcriptase (NEB, M0253S) by following the manual of the manufacturer. qRT-PCR was carried out with specific primer sets (listed in Supplementary Table 10) using SensiFAST™ SYBR® No-ROX Kit (BIOLINE, BIO-98005) and CFX Connect Real-Time PCR Detection System (BIO-RAD) with the following conditions: initial denaturation at 95 °C for 2 minutes, and 40 cycles of denaturation at 95 °C for 5 seconds, annealing at 55 °C for 10 seconds and extension at 72 °C for 15 seconds. cDNA concentrations were normalized with Ef1-α (elongation factor 1-alpha) housekeeping gene.

### Generation of reference database

A non-redundant set of 119,347 Axolotl proteins is computed from 181,431 RNA transcripts (Nowoshilow et al. 2018) in multiple steps. First, the pairwise identities of transcripts are computed through all-against-all BLAST (Altschul et al. 1990) searches using the transcripts database. Next, transcripts with 100% pairwise sequence identity are clustered together and the longest transcript for each cluster is selected as the representative. Finally, for each representative, the longest open reading frame among all six reading frames is translated into a protein sequence to generate the protein set. Furthermore, in order to support functional annotation of these Axolotl proteins, the most similar mouse protein orthologs for Axolotl proteins were identified by running BLAST searches against the mouse proteins from NCBI nr database (Barrett et al. 2015).

### Principal Component Analysis, Heatmap Generation and Clustering

Principal Component Analysis (PCA) was computed using R Stats (R Core Team 2018) package’s prcomp method and was plotted using R factoextra (Kassambara et al. 2017) package’s fviz_pca_ind method. DE proteins in metamorphic and neotenic samples for Day 0, Day 1, Day 4, and Day 7 amputation timepoints were visualized using “pheatmap” package in R (version: 3.5.1) (Team 2018; Kolde 2019). The clustering and heatmap construction were performed on tho se DEGs that pass our significance criteria; an adjusted P < 0.01 and fold changes (FC) for day1/day0, day4/day0, and day7/day0 being either > 2 or < −2). The distance measure that was used in clustering the samples was “Manhattan” distance. The genes were hierarchically clustered using “Complete Linkage” clustering method. For DE proteins in metamorphic day7 versus neotenic day7 samples, “Correlation” distance and “Average Linkage” was used for clustering the heatmap and visualization by using ClustVis online tool.

### GO and KEGG enrichment analyses

DE proteins of metamorphic day7/day0, neotenic day7/day0 and metamorphic day7/ neotenic day7 samples (significance thresholds: adjusted P < 0.01 and |FC| > 2) were tested for identifying KEGG pathways and Gene Ontology annotations spanning the three categories; biological processes, cellular components, and molecular functions using Bioconductor’s “clusterProfiler” package in R (Gentleman et al. 2004; Yu et al. 2012; Huber et al. 2015; Team 2018). Some important parameters of the enrichment test were set to a p-value and q-value cutoffs of 0.05 each, “org.Mm.eg.db” (mouse) as the organism database, the whole mouse database as the background (universe) gene list, and Benjamini & Hochberg (BH) as the adjusted P value method.

### Visualization of GO and KEGG terms

Heatmap-like plots and a dot plot were implemented for functional classification of KEGG enrichment results using Bioconductor’s “clusterProfiler” package in R. The latter package was also used to visualize metamorphic day7/ neotenic day7 enriched KEGG pathways by a dot plot. Pathway-based data integration and visualization of the “complement and coagulation cascades” KEGG term was generated using “pathview” package in R (Gentleman et al. 2004; Luo and Brouwer 2013; Huber et al. 2015; Team 2018). Bar plots were used to visualize the top 10 terms of each of the three GO annotation categories (biological processes, cellular components, and molecular functions) using “ggplot2” package in R (Wickham 2016, 2017; Team 2018).

### DE proteins screening and correlation testing

The R “ggplot2” package (Wickham 2016, 2017; Team 2018) was used to generate a scatter plot for the correlation between the fold changes of metamorphic day7/day0 DE proteins and those of Neotenic D7/D0 (significance thresholds: adjusted P < 0.01 and |FC| > 2). Spearman correlation test was used to calculate the correlation coefficient for the latter comparison using base R.

#### Protein-protein interaction (PPI) network analysis

Physical and functional interactions among the DE proteins in metamorphic day7/ neotenic day7 comparison were predicted using the Search Tool for the Retrieval of Interacting Genes (STRING; https://string-db.org/) online database (Szklarczyk et al. 2017). Interactions between up-regulated and down-regulated proteins were visualized using Cytoscape (version: 3.6.0) (Shannon et al. 2003).

